# STDrug enables spatially informed personalized drug repurposing from spatial transcriptomics

**DOI:** 10.64898/2026.04.03.715101

**Authors:** Yiwen Yang, Thatchayut Unjitwattana, Shu Zhou, Suguru Kadomoto, Xiaotong Yang, Tianao Chen, Abdullah Karaaslanli, Yuheng Du, Wanxing Zhang, Haodong Liang, Xuhui Guo, Evan T. Keller, Lana X. Garmire

**Author notes:** These authors contribute equally to this work (Yiwen Yang, Thatchayut Unjitwattana).

## Abstract

Drug repurposing offers a scalable route to accelerate therapeutic discovery, yet existing approaches based on single-cell RNA sequencing (scRNA-seq) often overlook spatial tissue context, limiting their ability to capture microenvironment-dependent drug responses. Here we present **STDrug**, a spatially informed computational framework that integrates spatial transcriptomics, graph-based modeling, and multimodal learning to enable patient-specific therapeutic prioritization. STDrug identifies and aligns disease and control spatial domains using graph convolutional networks and coherent point drift, and prioritizes candidate drugs through an integrative scoring scheme combining tumor-reversible gene signatures, perturbation-based reversal scores, and knowledge-guided gene weighting within a machine learning framework. By modeling spatial domain interactions alongside predicted drug efficacy and toxicity, STDrug generates robust patient-level drug scores. Across hepatocellular carcinoma and prostate cancer datasets, STDrug outperforms existing single-cell and spatial transcriptomics–based drug repurposing methods, achieving signficantly improved predictive accuracy (AUCs=0.81-0.82) across patients. Validation using large-scale electronic health records and in vitro assays further supports the translational relevance of top-ranked candidates. Taking together, STDrug establishes a generalizable framework for incorporating spatial omics into therapeutic discovery, advancing spatially informed and personalized drug repurposing.

## Introduction

Drug discovery remains a time-consuming and costly process, with most candidates failing before reaching the clinic^1^. Drug repurposing – identifying new therapeutic indications for approved or investigational compounds – offers a faster and less risky alternative by leveraging existing pharmacological and safety data^2^. Unlike de novo discovery, which relies on target identification and compound screening, drug repurposing strategies exploit diverse molecular, clinical, and computational resources to reposition known drugs toward unmet clinical needs.

Computation has become central to drug repurposing. Based on analytical approaches, it can be classified as network-based methods^3,4^, machine learning and deep learning models^5,6^ etc. Depending on the input data, drug repurposing can be categorized by drug-centric, disease/target-centric, or multi-modal approaches. Among them, omics-driven approaches, in particular transcriptomic-based approaches reversing disease-associated molecular signatures, have shown success^7,8^. However, the earlier transcriptomics-based approach used bulk gene expression, overlooking the cellular heterogeneity within the tissue that may significantly bias the disease associated signatures^9^.

To address this issue, some recent work was developed for drug repurposing, using single-cell RNA-seq (scRNA-Seq) data as the input. For example, Beyondcell^10^ is a framework that utilizes scRNA-seq data to identify tumor subpopulations with distinct drug response signatures. It prioritizes cancer-specific therapeutic candidates based on enrichment of drug sensitivity profiles. DrugReSC^11^ integrated single-cell transcriptional landscapes with compound-induced perturbation profiles to systematically identify repurposable drugs tailored to cellular phenotypes. We developed ASGARD, a framework that accounts for cell-type heterogeneity in scRNA-seq data to derive the comprehensive drug score holistically, by searching the candidate prepurposeful drugs through the L1000 pharmacogenomics database^12^. However, a major limitation of scRNA-Seq data is the lack of spatial context and organization of the single cells, which may include important clues on drug treatment effect^13^.

To address this issue, we here propose STDrug, a new computational method dedicated to using spatial transcriptomics data to aid drug repurposing. STDrug exploits spatial information specifically in identifying spatial domain pairs between the disease and healthy tissue. It proposes potential drugs and compounds from large-scale pharmacogenomics perturbation data, using a machine-learning method that learns the importance of the disease-associated genes by referring to the ground truth obtained from pre-trained large language models (LLMs). It offers realistic estimation for repurposeful drugs by considering drug efficacy, toxicity, and potential side effects. In short, STDrug aims to bridge a major gap currently, in bringing the cutting-edge technology of spatial transcriptomics towards guiding therapeutic treatment decision-making.

## Results

### The architecture of STDrug

STDrug comprises three major steps, including 1) data preprocessing and integration, 2) spatial domain identification, and 3) drug repurposing **(Fig. 1)**. STDrug first takes the spatial transcriptomic data from paired diseased and normal tissues, ideally from the same patient, as the input. It then performs a batch effect correction and sample alignment to reduce variability between patients. Inspired by SpaGCN^14^, we used batch-corrected Harmony embeddings rather than PCA to combine spatial coordinates and build a graph convolutional network (GCN) that learns spatially-aware low-dimensional embeddings. Different from SpaGCN, these embeddings are uniquely adjusted by a disease-control balanced clustering algorithm to identify paired spatial domains between the diseased and normal spatial transcriptomics slides for a patient.

**Figure 1:**
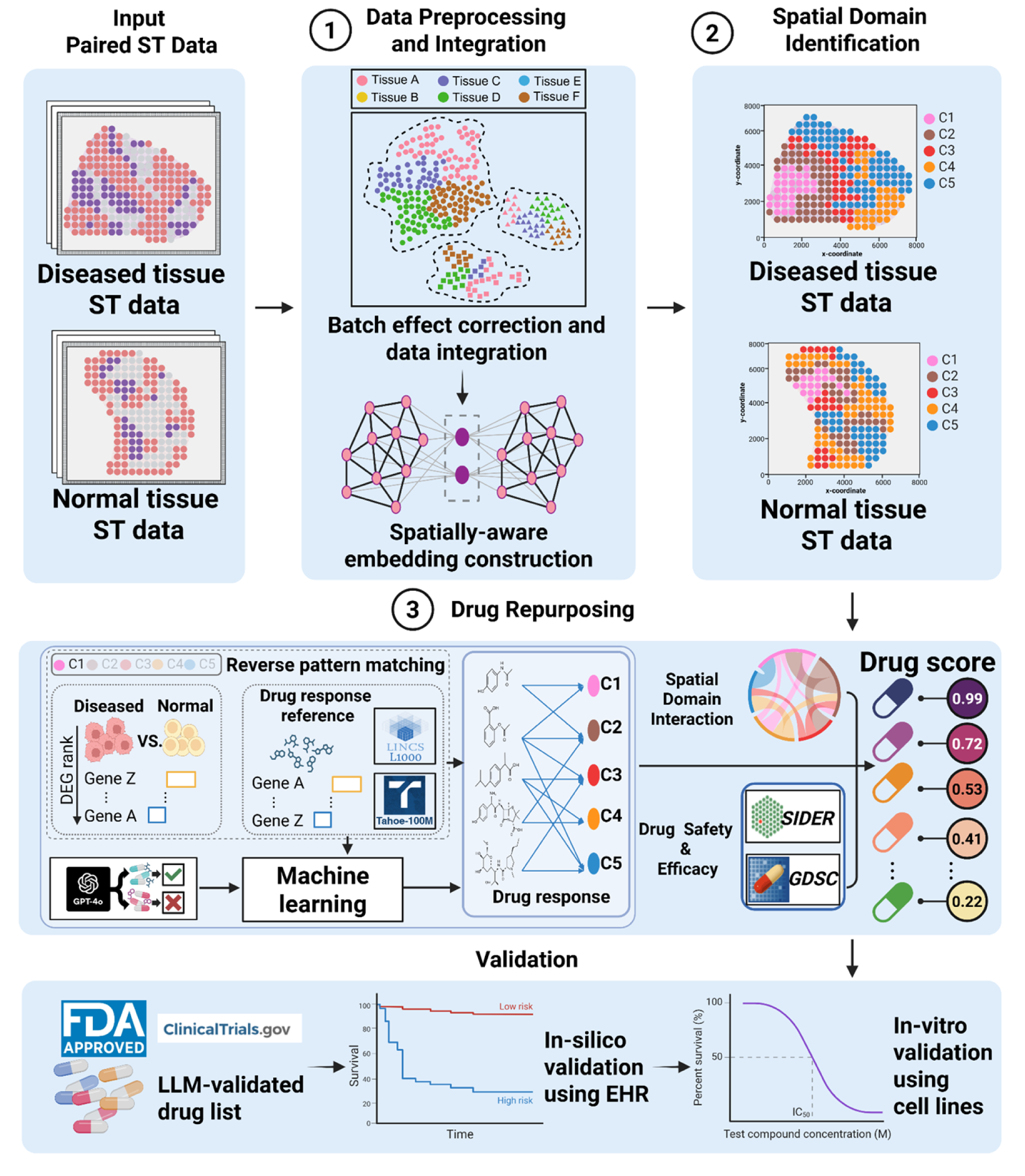
Illustration of STDrug architecture. STDrug utilizes paired diseased and normal tissues as the input. Data preprocessing, batch correction, and sample alignment are performed initially, before spatially-aware embeddings are constructed to identify corresponding spatial domain pairs between the diseased and normal conditions. These paired spatial domains then serve as inputs for the drug repurposing module, which identifies potentially reversible disease-associated genes determined as differentially expressed genes (DEGs) between spatial domains in diseased vs normal conditions. These disease-associated genes are used as the “queries” to search candidate drugs that may reverse these genes expression in the pharmacogenomics perturbation data from the L1000 and Tahoe-100M single-cell RNA-seq datasets. To prioritize key reversible disease-associated genes, STDrug assigns them weights learned from a machine learning model, where the gene-disease association was obtained from a pre-trained GPT-4o model on existing literature. Additionally, STDrug accounts for spatial domain interactions, drug side effect profiles from SIDER and sensitivity data from GDSC in calculating the comprehensive drug score for each candidate drug. The top-ranked drugs are then validated using a combination of evidence, including existing evidence from literature and clinical trials, in-silico validation using large-scale clinical data, and in-vitro validation from cell line experiments.

The resulting paired spatial domains serve as the foundation for the drug repurposing module, which prioritizes therapeutic candidates that maximally reverse the disease-associated gene expression changes from the pharmacogenomics perturbation databases (eg. L1000 and Tahoe-100M). The drug repurposing module utilizes potential drug information obtained from GPT-4o^17^, a pre-trained LLM by OpenAI, along with a ML method to assign weights to reversible disease-associated genes. Doing so prioritizes those drugs most likely to play key roles. In addition to reversible disease-associated genes, the drug repurposing also considers spatial domain interactions, as well as drug side effect information from the Side Effect Resource (SIDER) database (version 4.1)^19^ and drug efficacy from Genomics of Drug Sensitivity in Cancer (GDSC) database^20^. The resulting comprehensive drug score is calculated by integrating the multiple factors stated above. The most promising candidate drugs are those with top-ranking drug scores at the patient-specific level.

To demonstrate the utility of STDrug, we validate predicted top drug results using a combination of (1) empirical evidence from literature, clinical trials, and LLM-based validation, (2) in-silico validation using the real-world clinical data, and (3) in-vitro validation using multiple cell line experiments. Together, this study presents a spatially-informed, patient-specific, end-to-end drug repurposing framework that systematically integrates spatial transcriptomics, machine learning, and pharmacogenomics drug perturbation data, rigorously validated by real-world clinical and experimental evidence.

### Spatial domain identification and benchmarking

To evaluate the clustering results generated by STDrug, we benchmarked STDrug against four popular spatial domain identification tools, including STAGATE^24^, STAligner^25^, SPACEL^26^, and GraphST^27^. Due to the lack of ground truth for spatial domains from paired tumor and normal tissues which are crucial for the drug repurposing module of STDrug, we simulated paired tumor and adjacent normal dataset from real HCC data, where each spatial domain is occupied by a cells of the same type whose gene expression values are based on its scRNA-Seq datadisribution **(Fig. 2a)**. STDrug yielded the best averaged values of normalized mutual information (NMI) and adjusted rand index (ARI) that are significantly better than other methods as shown in **Fig. 2b** (NMI = 0.93, ARI = 0.94). STAGATES yielded the 2^nd^ best ARI and NMI (NMI = 0.78, ARI = 0.79), followed by STAligner (NMI = 0.76, ARI = 0.75). Comparatively, SPACEL (NMI = 0.36, ARI = 0.30), and GraphST (NMI = 0.24, ARI = 0.19) have the lowest accuracies (**Fig. 2b)**. Detailed investigation on the slide level of simulated spatial transcriptomes of each patient shows that the spatial domains identified by STDrug is consistently better aligned with the spatial domain ground truth, compared to other spatial domain identification methods **(Fig. 2c)**. These results highlight the robustness and accuracy of STDrug in spatial domain identification which relies on precise contextual mapping between tumor and normal tissue, reinforcing its suitability for drug repurposing by STDrug.

**Figure 2:**
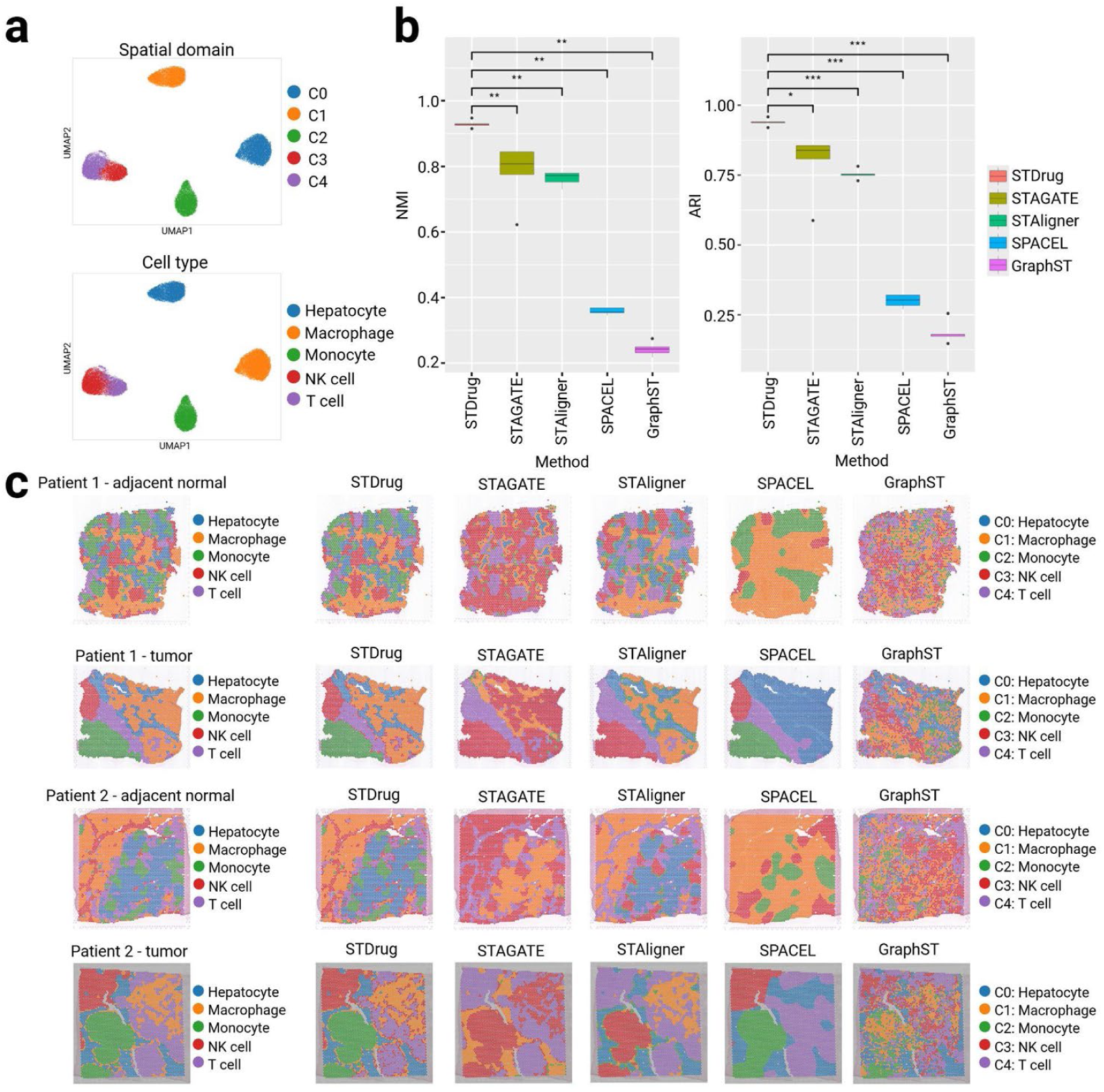
Spatial domain identification and benchmarking. a, UMAP plots of STDrug-identified domains (top) and cell types in the domains (bottom) in the simulated ST dataset. b, NMI and ARI of different spatial domain identification methods. Asterisks represent the significance of the statistical test (p-value), *** = 0.001, ** = 0.01, * = 0.05. c, Visualization of spatial domains estimated by different spatial domain identification methods on simulated spatial transcriptomics data from tumor and tumor-adjacent normal tissues from patients 1 and 2.

### Drug repurposing and benchmarking

Since interactions between spatial domains may also impact cellular processes within tissues^28^, STDrug also exploits domain-domain interactions (DDI) for drug score calculation. It computes the ratio of interaction strength between each pair of spatial domains by CellChat v2^29^. Domains exhibiting larger differences in DDI in diseased vs control conditions are considered more influential, when other parameters are equal. By integrating the proportions among all spatial domains, their DDIs, as well as the therapeutic connectivity FDR, a patient-level comprehensive therapeutic score is computed. To further take into account both drug efficacy and safety, STDrug can optionally include drug efficacy and safety scores in the comprehensive drug score, by leveraging the drug sensitivity from the GDSC database^20^ and drug side effects information from the SIDER database^19^. To enable comparison among patients, the drug score is rank-scaled between 0 and 1.

We tested STDrug by predicting potential drugs for hepatocellular carcinoma (HCC) and prostate cancer (PCa) treatments. The HCC data are taken from four patients with paired tumor and adjacent normal tissues^30^ and the PCa data are from two patients with paired tumor and adjacent normal tissue spatial transcriptomics data, where the non-tumor regions were annotated by pathologists^31^. Despite the existence of drug response prediction methods, the field currently generally lacks dedicated spatial-transcriptomic-based drug repurposing methods in which drug selection and ranking is the objective, except Beyondcell^10^ which was extended to spatial transcriptomics data We also compared with ASGARD^12^ in which scRNA-Seq is the input.We used the area under the ROC curve (AUC) to evaluate the prediction accuracy, where the “proxy truth data” of drug-disease association are obtained from clinical trials, and a pre-trained GPT-4o model using a hold-out set of literatures independent from those used to train the XGBoost model^17^. Note the hold-out set of literature from GPT-4o search is separate from those used to train the XGBoost model.

For HCC data, STDrug achieves an AUC of 0.81, significantly outperforming both ASGARD (AUC = 0.61) and Beyondcell (AUC = 0.59) in identifying potential drugs (**Fig. 3a)**. The significance of potential top drugs across four HCC patients are shown in **Fig. 3c**. STDrug consistently identifies promising drug candidates across HCC patients, with bosutinib and sorafenib as top candidates in four patients. Sorafenib is the first-line chemotreatment for advanced HCC patients, acting by suppression of angiogenesis and tumor proliferation^32,33^. The top ranking of Sorafenib confirms the strong potential of STDrug in drug repurposing prediction for HCC. Bosutinib, an FDA-approved tyrosine kinase inhibitor for Philadelphia chromosome-positive chronic myeloid leukemia, has been reported to exhibit anti-tumor activity in preclinical models of hepatocellular carcinoma^34^. STdrug also predicted repurposeful drug used to treat other cancers such as daunorubicin (for AML and ALL), dactinomycin (for rare pediatric cancers such as Ewing sarcoma), trametinib (melanoma), panobinostat (multiple myeloma), and epirubicin (breast, colon and gastric cancers). STDrug additionally identified a diverse array of therapeutic candidates in at least one patient – including non-drug supplements niacin (vitamin B3) and caffeine, and medications for treating cardiovascular diseases including atorvastatin, digoxin, and losartan. Niacin has been reported to suppress liver tumor growth by enhancing antitumor immunity^35^. Caffeine has been shown to HCC tumor suppressing functions, ranging from cell lines^36^, mouse models^37,38^, to large-scale epidemiological studies^39–42^. Atorvastatin, commercial name as Lipitor, is used to prevent heart attack and stroke clinically. It has been clinically shown to improve HCC prognosis^43,44^. Similarly, digoxin (Lanoxin), a cardiac glycoside commonly used for heart failure and atrial fibrillation, has been reported to suppress angiogenic and tumor growth in HCC^45^.

**Figure 3:**
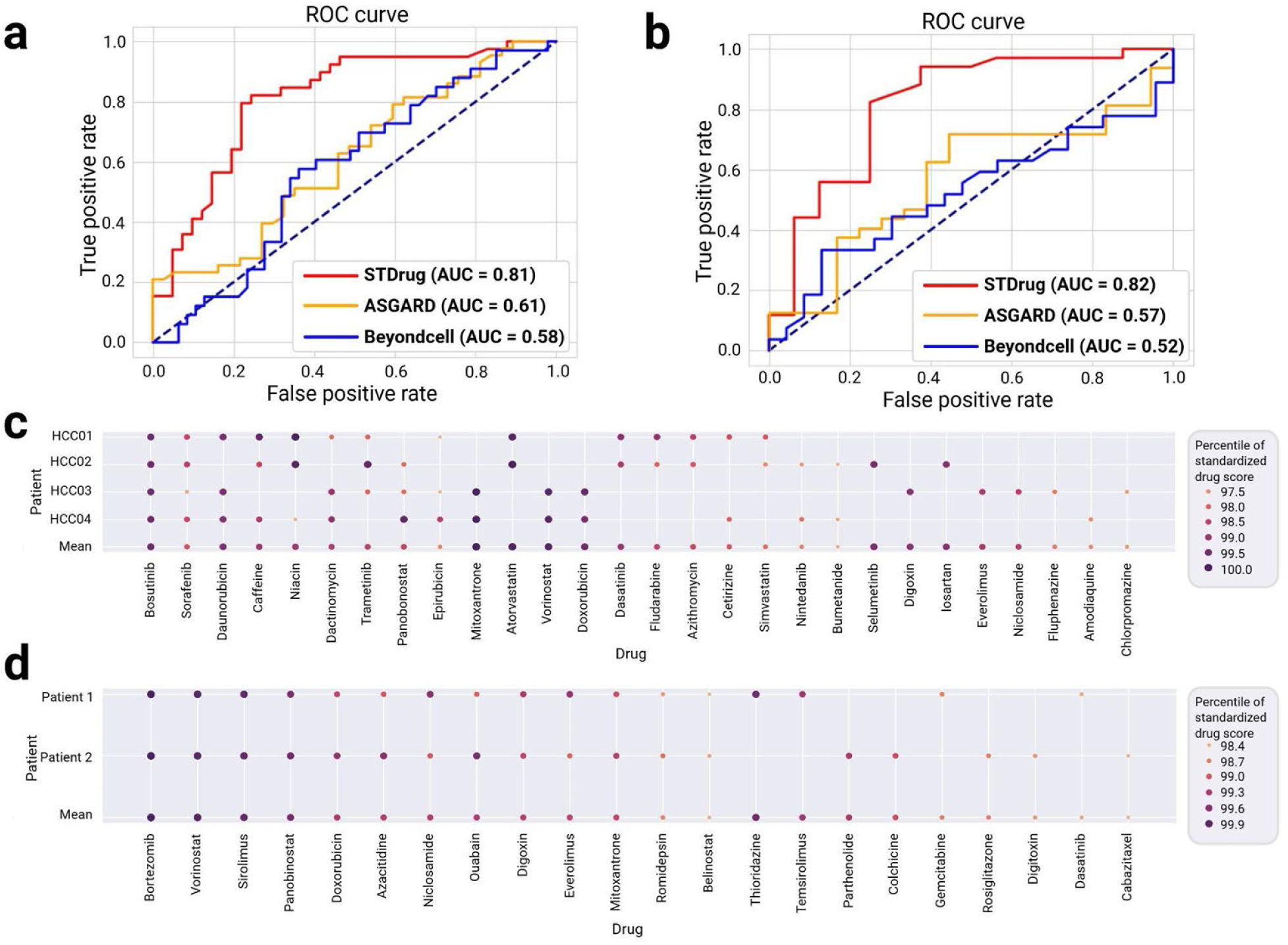
Drug repurposing and benchmarking. a-b, Evaluation of drug repurposing tools on HCC tissues (a) and PCa tissues (b). For each patient, only top 20 drugs are selected and ranked by normalized drug score percentages (%). For the averaged ranking across patients, drugs are ordered in descending order based on their frequency of occurrence among patients, followed by the mean percentile of normalized drug scores among only the patients who have the drugs enlisted as top 20. c, STDrug-identified top candidate drugs and compounds for HCC patients. d, STDrug-identified top candidate drugs and compounds for PCa patients.

For PCa, STDrug also shows superior performance with an AUC of 0.82, compared to ASGARD (AUC = 0.57) and Beyondcell (AUC = 0.48) (**Fig. 3b)**. It identifies 13 common drug and chemical compound candidates for PCa in the two patients **(Fig. 3d).** Among them, top five drugs have shown anti-prostate cancer functions, including bortezomib^46^, vorinostat^47^, sirolimus^48^, panobinostat^49^, and doxorubicin^50^. Interestingly, STDrug also prioritized ouabain and digoxin-agents traditionally used for heart conditions-as potential therapeutic candidates for prostate cancer.

### Validation of potential repurposeful drugs for HCC and PCa

We first conducted validations using the real-world patient clinical data from MarketScan (n=264 million patients). As patients diagnosed with cancers were seldomly given repurposeful new drug proposed here, we focused on the patients exposed to repurposeful drugs before cancer diagnosis. A subset of predicted repurposed drugs reached the patient size threshold (n=66) among the cancer of interest, after matching them with untreated controls by demographics and comorbidities. These drugs are atorvastatin, digoxin, and niacin for HCC, and digoxin, colchicine, pitavastatin, and sirolimus for PCa. As shown by Kaplan-Meier curves, atorvastatin, digoxin, and niacin all significantly delay the onset of HCC compared to those with no history of using these drugs **(Fig. 4a).** Digoxin, colchicine, pitavastatin, and sirolimus also exhibit notable delays in PCa development compared to the control group **(Fig. 4b)**. The Hazard Ratios (HR) of the Cox proportional hazards models are shown in **Fig. 4c-d**, further confirming the protective effects of these candidate drugs/compounds.

**Figure 4:**
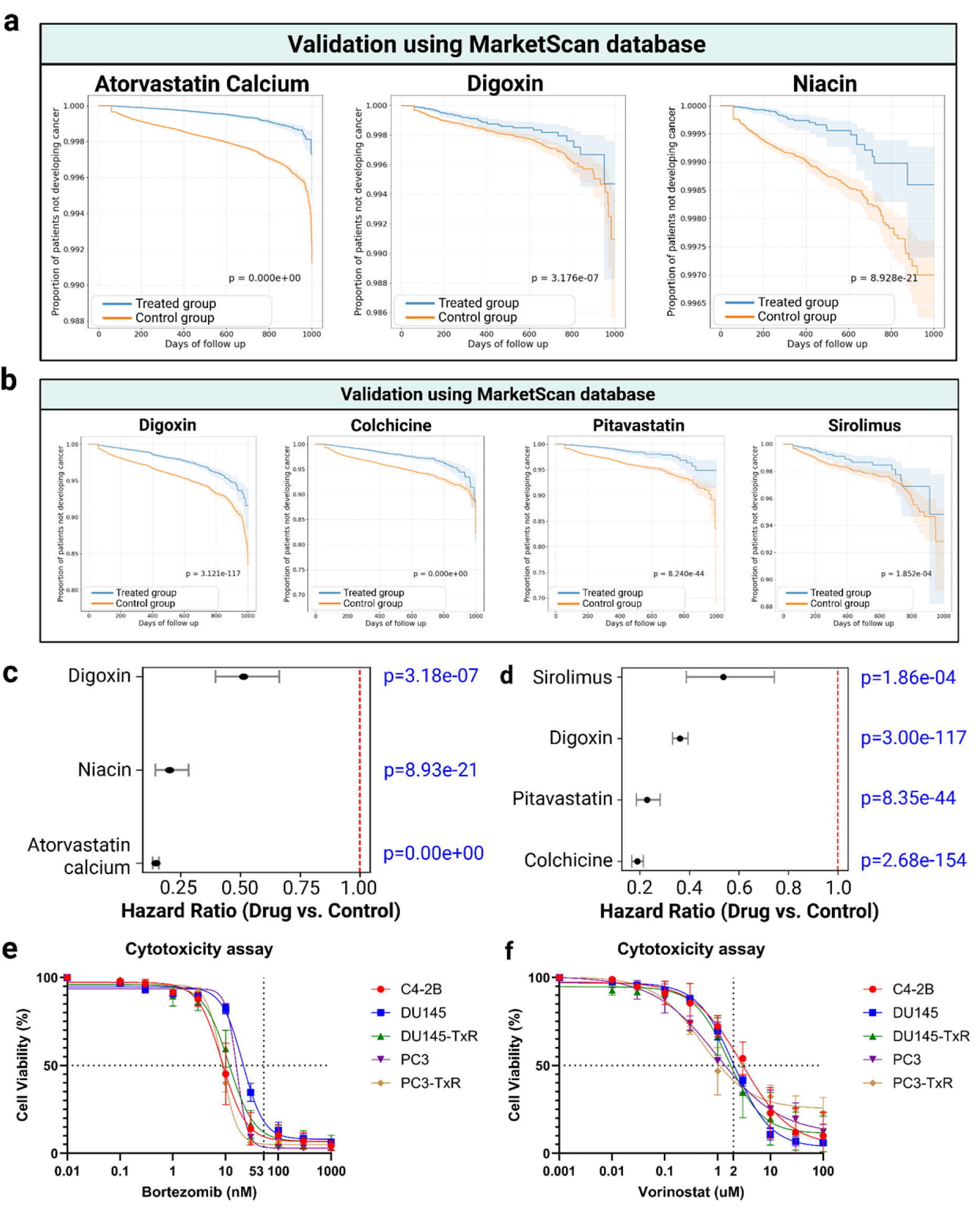
Real-world and in-vitro validation of top candidate drugs. a, Real-world clinical data validation of potential HCC drugs using Marketscan database. b, Real-world clinical data validation of potential Pca drugs from MarketScan database. c-d, Hazard Ratios (HR) of Cox proportional hazards models for the drug/compound candidates for (c) HCC and (d) PCa from Marketscan database. e-f, In-vitro validation of (e) bortezomib and (f) vorinostat in prostate cancer cell lines. The dotted horizontal line marks 50% reduction in cell viability, and the dotted vertical line represents clinical blood concentration of the drug.

Additionally, we conducted in-vitro validation of the repurposed drug candidates for PCa by cell viability assays. As complement, we evaluated the effects of bortezomib and vorinostat, the candidates which lack sufficient clinical records among PCa patients. We tested five prostate cancer cell lines, including C4-2B, DU145, DU145-TxR, PC3, and PC3-TxR. **(Fig. 4e-f)**. Bortezomib, a protease inhibitor used in multiple myeloma^51^, was effective against all prostate cancer cell lines (IC50: C4-2B = 8.57 nM, DU145 = 21.2 nM, DU145-TxR = 11.4 nM, PC3 = 16.3 nM, PC3-TxR = 8.55 nM; clinical blood concentration = 53 nM^52^). Vorinostat, a histone deacetylase (HDAC) inhibitor approved for the treatment of lymphoma^53^, also shows selective efficacy against DU145-TxR, PC3 and PC3-TxR prostate cancer cell lines (IC50: DU145-TxR = 1.59 uM, PC3 = 0.963 uM, PC3-TxR = 0.508 uM; clinical blood concentration = 2uM^54^). Taxane-resistant cell lines (PC3-TxR and DU145-TxR) exhibit slightly higher sensitivity to vorinostat compared to their parental counterparts (PC3 and DU145). These findings highlight the potential of bortezomib and vorinostat for prostate cancer treatment, especially in the treatment-resistant prostate cancer subtypes.

### Bioinformatics analysis of the effect of HCC candidate drugs

To interpret how candidate drugs may affect the cells in HCC ST data, we first performed analysis of spatial domains from HCC tumor and adjacent normal tissues (**Fig. 5a, b).** On each spatial domain, we conducted cell type deconvolution using cell2location^55^ (**Fig 5c)**. As a result, five related but transitional spatial domains are detected optimally (**Fig 5a**), characterized as a proliferative adaptive domain C1, a fibrotic immune-evasive domain C2, a transitioning immunoactive domain C3, an immune-limited adaptive domain C4, and an immunosurveillance-disrupted domain C5.

**Figure 5:**
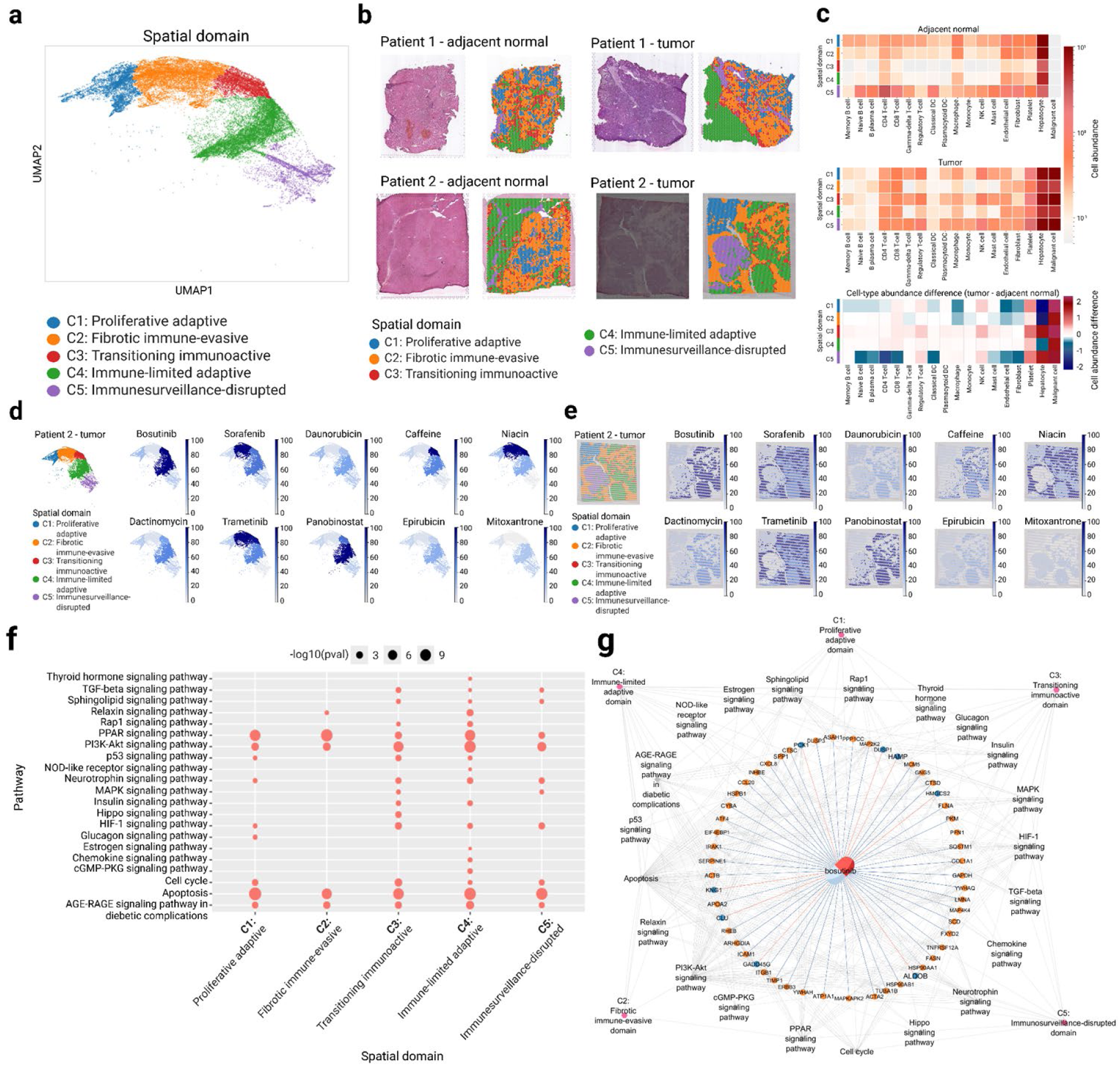
HCC functional downstream analysis. a, UMAP plot of HCC tissues colored by STDrug domains. b, Visualization of STDrug domains on tissues from patients 1 and 2. c, cell type deconvolution of HCC tumor and adjacent normal tissues. d-e, drug score visualization of top 10 drugs from Figure 3d on (d) UMAP plot and (e) spatial plot of patient 2. f. Target pathways of bosutinib in patient 2. g, The top drug candidate, bosutinib, its target genes, pathways, and domains. Orange node: up-regulated gene (logFC>1 and adjusted *P*-value < 0.05). Blue node: down-regulated gene (logFC < −1 and adjusted *P*-value < 0.05). Orange solid edge: drug stimulates gene expression. Blue solid edge: drug inhibits gene expression. The width of the edge is proportional to the strength of the drug effect. Gray dotted edge: gene belonging to a pathway. Gray backward slash: pathway significant in a corresponding domain.

To observe drug effects in these spatial domains, we visualized the drug scores stratified by spatial domains. As an example, **Fig. 5d-e** shows 10 top drugs on patient 2, by UMAP and 2D-geolocation plot respectively. The main targeted regions by candidate drugs vary and can be roughly divided into 2 classes. Class 1 includes sorafenib, niacin, and trametinib that main act on spatial domains 2, 3 and 4, particularly domain 2 which is enriched with hepatocytes but relatively low in fibroblasts and immune cells. Others (eg. bosutinib, caffeine and panobinostat) mainly act on spatial domains 3 and 4. In particular, caffeine exerts the most significant effect on domain 3- the transitioning immunoactive domain.

To explore the potential molecular mechanism of the top drug candidate bosutinib, we analyzed enriched pathways of the putative target genes (gene that have reversed expression under bosutinib, compared to the HCC condition) across the five spatial domains **(Fig. 5f)**. While a few pathways are activated across all spatial domains, such as PPAR, PI3K-Akt and apoptosis pathways, pathway dys-regulation is much more prevalent in spatial domains C3 and C4 as compared to C1 and C2. This is consistent with that C3 and C4 are the main spatial.domains affected by bosutinib. In domain C3 (the transitioning immunoactive domain) MAPK signaling is enriched (**Fig 5f**), supported by the evidence that overexpressed pro-oncogenic MAP2K2 (MEK2) and MAP4K4 in the MAPK/ERK cascade are predicted to be down-regulated by bosutinib indicated by blue edge (**Fig. 5g**). As bosutinib is an Src/Abl tyrosine kinase inhibitor, ITGB1, a gene in the canonical Src-associated signaling pathway, is predicted to be down-regulated as expected. In addition, multiple cytoskeleton-associated genes that are part of Src-regulated adhesion and motility programs are predicted to be down-regulated by bosutinib, including FLNA and PFN1 in the actin-remodeling components, and ACTB and ACTA2 in the actin filaments. Additionally, molecular chaperones HSP90AA1 and HSP90AB1, which was linked to stability of the Src signaling axis^56,57^, are also predicted to be downregulated by bosutinib, indicated by blue edges in **Fig. 5g**. Overall, Src-regulated biological functions show repression, being the targets of bosutinib.

### Bioinformatics analysis of the effect of PCa candidate drugs

We conducted similar spatial domain analysis in the PCa patients. Interestingly, STDrug again identifies five optimally distinct domains after integrating the tumor and normal tissues (**Fig 6a, 6b**). We classify them as the vascular remodeling domain C1, fibrovascular stromal domain C2, adaptive immune-regulated domain C3, metabolic and immune-responsive domain C4, and epithelial-driven tumor expansion domain C5. As expected, the tumor samples have much higher presence of C5 but much lower C2 domains (**Fig 6b**). The cell-type deconvulation results show that tumor has attracted significantly more naive T-cell and B-cells to the site, however is reduced in fibrobrasts and pericytes (**Fig 6c**). Furthermore, we investigated the effect on the spatial domains from top common drugs, using patient 1 as an example **(Fig. 6d-e)**. Most drug candidates act on multiple domains. Despite significant spatial domain-level differences in the drug scores among these drugs, C3 domain (adaptive immune-regulated domain) appears to be a common targeted spatial domain. C3 domain is characterized by significant increase of B-cell and naive T-helper cell population but largest decrease of fibroblasts in tumor samples (**Fig 6c)**. C1 domain (vascular remodeling domain) is the other common, but less-strongly targeted spatial domain. Corroborating with **Fig 3e**, Bortezomib and Vorinostat, the top 2 most significant drugs, exert the strong effects on all domains except C5 (**Fig 6e**). Sirolimus targets C3 relatively the strongest, followed by panobinostat, niclosamide, everolimus, ouabain, azacitidine, and doxorubicin in the descending order of the drug score over C3.

**Figure 6:**
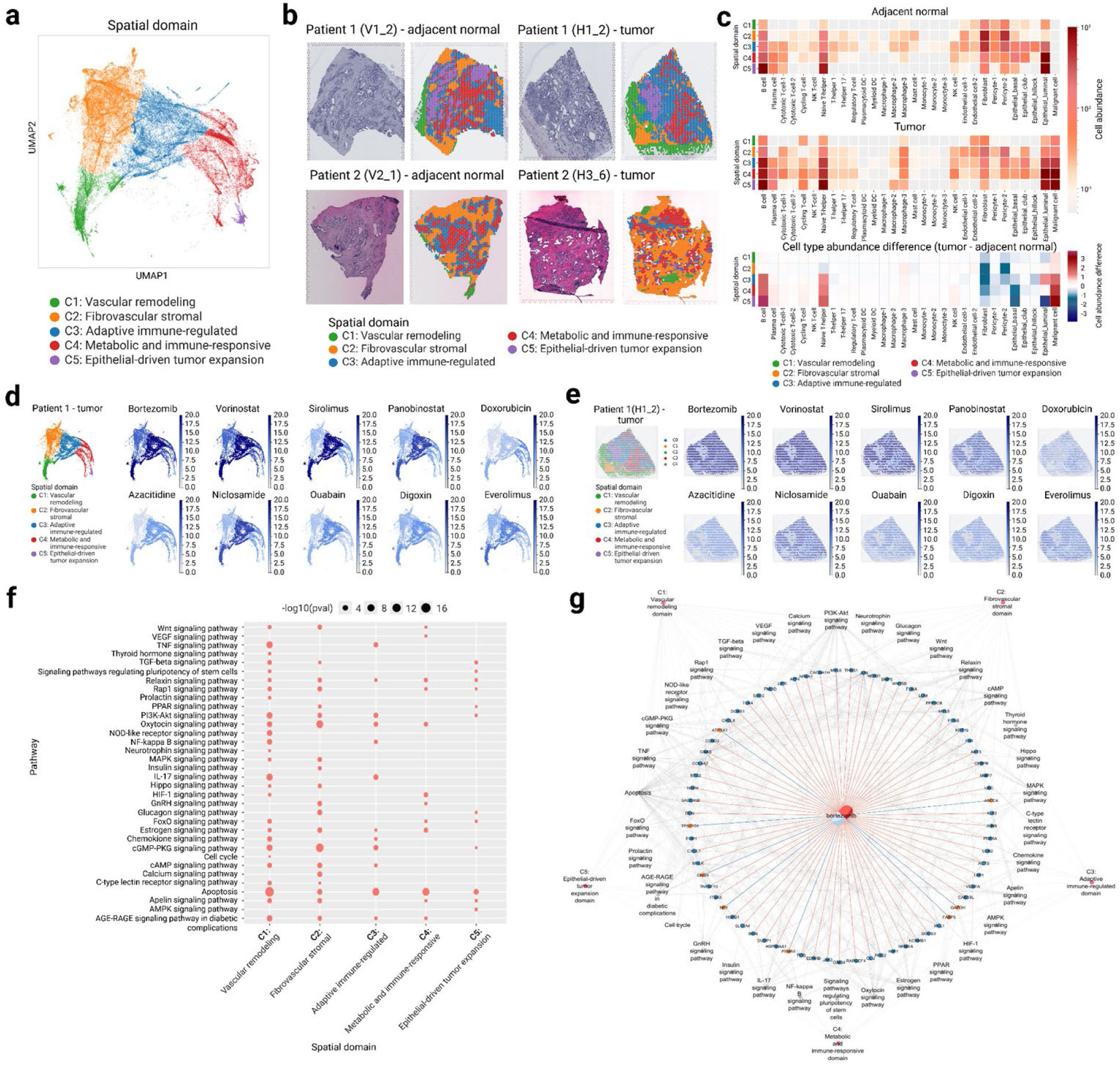
Prostate cancer functional downstream analysis. a, UMAP plot of prostate cancer tissues colored by STDrug domains. b, Visualization of STDrug domains on tissues from patients 1 and 2. c, cell type deconvolution of prostate tumor and adjacent normal tissues. d-e, drug score visualization on (d) UMAP plot and (e) spatial plot of patient 1. f, Target pathways of bortezomib in patient 1. g, The top drug candidate, bortezomib, its target genes, pathways, and domains. Orange node: up-regulated gene (logFC>1 and adjusted *P*-value < 0.05). Blue node: down-regulated gene (logFC < −1 and adjusted *P*-value < 0.05). Orange solid edge: drug stimulates gene expression. Blue solid edge: drug inhibits gene expression. The width of the edge is proportional to the strength of the drug effect. Gray dotted edge: gene belonging to a pathway. Gray backward slash: pathway significant in a corresponding domain.

To further elucidate the molecular mechanisms underlying the top candidate drug bortezomib, we analyzed the genes affected as well as their enriched pathways across the spatial domains. **(Fig. 6f,g)**. Corresponding to the domain level drug impact, the altered pathways in C5 are the least and weakest. Some most significant pathways include apoptosis in spatial domains C1-C4, which is a known consequential effect of bortezomib; cGMP-PKG signaling pathway and PI3K-Akt signaling pathway in domains C1-C3, and TNF and IL-17 signaling pathways in domains C1 and C3. Some of the genes previously known to be affected by bortezomib are among the putative target genes including the direct drug target PSMB5 and downstream genes TNFSF10, DCN, and WIF1 **(Fig. 6g)**. PSMB5 level is upregulated in prostate cancer (red node) and repressed by Bortezomib as indicated by the blue edge (**Fig. 6g)**. TNFSF10, encoding the pro-apoptotic ligand TRAIL, is downregulated in the tumor (blue node) and predicted to be restored by bortezomib (red edge). Two prostate cancer tumor suppressors, DCN and WIF1, are also downregulated and predicted to be indirectly rescued by bortezomib (red edge).

## Discussion

We present STDrug (Spatial Transcriptomics aided Drug Repurposing), a computational framework that enhances drug repurposing by leveraging spatial transcriptomic data. Beyond data preprocessing and integration, STDrug is characterized by two unique and innovative modules: the spatial domain identification module and the drug repurposing module. The spatial domain identification module aligns matched spatial regions between diseased and normal tissues using a graph convolutional network (GCN) combined with coherent point drift (CPD) algorithm. The superior accuracy of spatial domain pair identification is confirmed by the benchmarking study with other tools, including STAGATE^24^, STAligner^25^, SPACEL^26^, and GraphST^27^.

We evaluated STDrug by various approaches, from literature/text-based validation, real-world EHR based validation, to in vitro experimental system. As no spatial drug-repurposing framework currently exists, we first compared STDrug to scRNA-seq-based approaches, including ASGARD^12^ and Beyondcell^10^, and evaluated by the literature and clinical trial data curated by Gen AI. STDrug yielded significantly higher AUCs in both hepatocellular carcinoma (HCC) and prostate cancer (PCa) patient data. Next, using real-world large-scale MarketScan clinical dataset for case-control studies, we confirmed the significant therapeutic effect of top drugs predicted by STDrug, including atorvastatin, digoxin and niacin for HCC, and digoxin, colchicine, pitavastatin and sirolimus for PCa. They were associated with delayed disease onset time and improved survival in propensity-adjusted Cox and Kaplan-Meier analyses. Lastly, we showed that in vitro cell culture across five PCa cell lines, including taxane-resistant PC3-TxR and DU145 models, confirmed potent cytotoxicity of bortezomib and vorinostat at clinically achievable concentrations^58–60^.

In HCC, STDrug prioritized bosutinib and sorafenib, with sorafenib serving as an internal positive control as a first-line therapy for advanced HCC. Bosutinib-predicted signatures included suppression of MAPK and Src-associated signaling (MAP2K2, MAP4K4, ITGB1), suppression of cytoskeletal regulators (FLNA, PFN1, ACTB, ACTA2), and reduced HSP90 chaperone activity (HSP90AA1, HSP90AB1), consistent with destabilization of oncogenic kinase networks and disruption of mitochondrial respiration^34^. Beyond kinase inhibitors, STDrug also identified several non-oncology agents with potential relevance to HCC. Niacin (vitamin B3) was associated with enhanced antitumor immunity through reduced immunosuppressive myeloid polarization and restoration of CD8 T-cell activity^35^, while caffeine has been linked to reduced HCC risk and suppression of proliferation via inhibition of adenosine signaling and PI3K-Akt activity^37,39–42^. Cardiovascular medicine including atorvastatin and digoxin were also highlighted, aligning with reported inhibition of MAPK signaling and hypoxia-inducible pathways, respectively^43–45,61^.

In PCa, STDrug prioritized bortezomib and vorinostat as top candidates, which were validated by the cell line experiments. Bortezomib, a proteasome inhibitor, exerts its antitumor activity by selectively and reversibly blocking the 26S proteasome – a multiprotein complex responsible for degrading ubiquitinated proteins involved in cell cycle regulation and apoptosis^62,63^. On the other hand, vorinostat, a histone deacetylase (HDAC) inhibitor, promotes histone hyperacetylation, leading to chromatin remodeling and the re-expression of silenced tumor suppressor genes, which induces cell cycle arrest at the G1 and G2/M checkpoints^64,65^. Further analyses indicated that bortezomib is associated with perturbation of PI3K-Akt, cGMP-PKG, TNF, and IL-17 signaling pathways in multiple spatial domains. This is consistent with coordinated modulation of survival, inflammatory, and androgen receptor-driven transcriptional programs^46,66–76^. These predictions align with established bortezomib mechanisms, including proteotoxic stress induction, G0/G1 arrest, and antiangiogenic effects in xenograft models^46,72^. Predictions on the downregulation of PSMB5, a direct proteasome target of bortezomib is consistent with established mechanisms of bortezomib on proteasome inhibition. Additionally, restoration of pro-apoptotic TNFSF10 (TRAIL) signaling and tumor suppressors DCN and WIF1, inferred from the spatial transcriptomics data by bortezomib treatment, also confirm its roles in the proteotoxic stress-induced apoptosis and tumor-suppressive signaling activation in prostate cancer^77–80^. Interestingly, STDrug also identified multiple endocrine pathways regulated by hormones as the potential targets of bortezomib, including relaxin, oxytocin, estrogen, and prolactin signaling, supporting the cross-talk with androgen receptor (AR)-driven transcriptional programs, which are central to prostate cancer biology^81^.

In summary, STDrug represents a comprehensive computational framework for drug repurposing using spatial transcriptomics information. By integrating spatial domain architecture, transcriptional reversibility, pharmacological constraints, and machine-learning and generative AI-based marker prioritization, it identifies clinically validated therapies. STDrug is expected to inform precision therapeutic discoveries, supported by mechanistic insight for repurposing from drugs and compounds.

## Methods

### Spatial transcriptomics (ST) dataset

To demonstrate the utility of STDrug, we applied it to two Visium ST datasets for two cancer types, HCC and PCa. The HCC Visium dataset consists of four patients, each with one tumor tissue and one matched adjacent normal tissue. The accession number for the raw sequencing data deposited in Genome Sequence Archive (GSA) is HRA000437 (https://ngdc.cncb.ac.cn/gsa-human/browse/HRA000437)^30^. The PCa Visium dataset consists of two patients, one with seven tumor tissues and two adjacent normal tissues, the other with four tumor tissues and 11 adjacent normal tissues. The spatial count matrices and high-resolution images are available on Mendeley Data (https://doi.org/10.17632/svw96g68dv.1)^31^.

### Processing of spatial transcriptomics data

We initiate the spatial transcriptomics data analysis by performing normalization and log transformation. The *normalize_total* function from Scanpy is used to scale the total counts per cell, followed by the application of the log1p function to stabilize the variance. Next, we reduce the data’s dimensionality through principal component analysis (PCA), where 50 principal components are computed using the *pp.pca* function. Following PCA, we apply the Harmony algorithm to correct for batch effects across different experimental conditions. This batch correction ensures the data are harmonized across patients, reducing unwanted technical variation while preserving biological signals.

### Functional downstream analysis of spatial transcriptomic data

Differential expression analysis was performed using the Seurat package (version 5.0.1)^82^. We performed GSEA using *gseKEGG()* in the package ClusterProfiler (4.8.2) with the gene set information from the Kyoto Encyclopedia of Genes and Genomes (KEGG) database^83,84^. The spatial domain interaction inference was performed using the package CellChat v2^29^. Cell-type deconvolution was performed using the Cell2location package^55^. Cell-type deconvolution of HCC tissues was performed using the integrated dataset of GSE189903^85^ and GSE149614^86^ as a reference. Cell-type deconvolution of prostate cancer tissues was performed using GSE181294^87^ as a reference. Cell-type dropout experiments were conducted by subtracting the average expression of the target cell type in the single-cell reference data from the spatial expression deconvolution results.

### Simulation of spatial transcriptomics data

Due to the lack of ground truth for spatial domains from paired diseased and normal tissues, we’ve developed a data simulation method to generate spatial transcriptomics (ST) data while ensuring paired spatial domains across conditions. For each tissue, we derived k spatial domains by jointly clustering cells based on their original gene expression profiles and the Euclidean distances between their spatial locations. Spatial domains between normal and tumor tissues were then matched, where a single cell-type cluster was assigned to a spatial domain (e.g. hepatocyte, macrophage, monocyte, NK cell, and T-cell). The cell type-specific gene expression distribution is derived from scRNA-seq reference data, following the Poisson distribution.

### Spatial Domain Identification

#### Spatially Aware Embedding

We construct a spatially aware representation by integrating batch-corrected low-dimensional gene expression embeddings with spatial neighborhood structure. Specifically, spatial graphs are built for each sample and incorporated into a unified graph learning framework, where expression features and spatial connectivity jointly inform representation learning. We employ a graph neural network architecture adapted for multi-sample inputs to capture shared and sample-specific spatial patterns. The model is optimized to learn latent embeddings that resolve spatial domains. These embeddings are subsequently used to identify conserved spatial structures across patients.

#### Disease-Control Balanced Clustering

The spatially aware embeddings were further processed using a nonlinear dimensionality reduction approach that preserves local neighborhood structure. To facilitate within-patient cross-condition comparison, embeddings derived from diseased and control tissues were aligned using a probabilistic non-rigid registration framework that enforces smooth spatial correspondences. Based on the aligned embeddings, we implemented a condition-balanced clustering strategy to jointly partition cells or spots across samples. Clustering was performed through iterative community detection under multiple resolution parameters, with constraints to ensure representation from both conditions and across patients within each cluster. An initial partition was obtained using a data-driven estimate of domain complexity, followed by iterative refinement steps that adjusted cluster granularity and reassigned underrepresented elements to achieve balanced partitions. The resulting clusters define spatial domains that are comparable across disease states and individuals.

### Drug Score Calculation

#### Construction of Integrated Drug Perturbation Reference

To construct a comprehensive reference for drug response, we integrated large-scale bulk and single-cell perturbation datasets into a unified framework. Single-cell perturbation profiles were aggregated into pseudo-bulk representations at the cell line level to ensure compatibility with bulk expression signatures. Expression profiles were standardized using within-sample normalization followed by robust scaling to enable cross-dataset comparability. The processed pseudo-bulk data were then harmonized with bulk perturbation data on a shared gene set and overlapping compound space. The resulting integrated reference comprises a large collection of perturbational signatures spanning multiple cell lines and tissue contexts.

#### DEG and CEG Z-score Calculation

Using the paired spatial domains identified from the STDrug clustering module, we compare the differentially expressed genes (DEGs) between tumor tissue and its adjacent normal tissue for each domain. Integrating gene expression profiles from the integrated drug perturbation reference, we calculate a standardized Z-score based on the probability of each gene being a concordantly expressed gene (CEG).

#### LLM-Enhanced Perturbed Gene Weights

To quantify the potential contribution of each CEG to the drug repurposing process, we introduce a perturbed gene weight by training the machine-learning classifier. To assist drug power labeling, we use gpt-4o (version 2024-08-06), a pre-trained large language model (LLM) from OpenAI to convert the base knowledge of the drug-disease relationship extracted from PubMed articles. We then calculate the feature gain for each gene.

#### Patient Therapeutic Drug Score

For each drug, a patient-specific therapeutic score was computed by integrating signals across spatial domains. This score accounts for domain composition, relative differences between conditions, and interaction patterns among spatial regions, resulting in a composite measure of predicted therapeutic relevance at the individual level. In addition, we incorporated complementary measures of drug efficacy and safety derived from external pharmacological and clinical knowledge sources. These measures were standardized and integrated to refine drug prioritization.

#### Repurposed Drug Ranking

The personalized drug score is a combination of the individual patient therapeutic score, drug efficacy score, and drug safety score. To facilitate comparison across different patients, we also provide a rank-based standardized drug score (percentile).

### Evaluation of potential repurposed drugs and benchmarking

Drug predictions from STDrug were evaluated using the area under the ROC curve (AUC), comparing predicted drugs against a reference set of known or validated drugs for each cancer type. To construct the validated drug reference list, we used FDA-approved cancer drug databases, clinical trial results from the NIH clinical trial registry (clinicaltrials.gov), DrugBank, and peer-reviewed literature from PubMed with supporting evidence from experimental or pharmacological studies. To benchmark STDrug against single-cell RNA-seq-based drug repurposing methods, including ASGARD and Beyondcell, we selected the top 80 HCC drugs and top 50 PCa drugs predicted by each method and compared their AUC scores against the validated drug reference list.

### In-silico validation of potential repurposed drugs

Merative MarketScan is a large private research database of de-identified US patient data, covering 264 million employees and their dependents from 2009 to 2022. It includes claims data, medical, drug, dental records, lab results, and hospital discharges, sourced from insurance providers, Blue Cross Blue Shield, and third-party administrators, offering insights into healthcare utilization and costs.

Our approach to validate STDrug repurposed drugs for the cancer types of interest using EHR data involved identifying treatment and control cohorts, and comparing their risk of developing cancer over a defined observation period. First, we designated patients who were administered or prescribed the drugs as the treated group. We then matched four patients from the untreated control group to each individual in the treated group based on birth year, sex, geographic location, and baseline comorbidities, including hypertension, type 2 diabetes (T2D), and coronary artery disease (CAD). Corresponding International Classification of Diseases (ICD) codes were used to identify the comorbidities diseases from individual records.

The observation period for the treated group was defined as the time between the first drug order and 60 days following the last drug order, provided there were no subsequent orders. For the control group, the observation period was matched to that of the treated group based on patient birth year, sex, geographic location and medical condition. For both the treated and control cohorts, we collected available medical records and identified individuals diagnosed with HCC or PCa based on ICD codes. Patients diagnosed within the observation period were marked as 1, while those who were not diagnosed were censored as 0.

We employed the Cox proportional hazards (CoxPH) model, stratified by propensity score, to analyze the relative risk of disease development between the treated and control cohorts. The propensity score was estimated using the same covariates applied in the case–control matching. Specifically, a logistic regression model was fitted to estimate each individual’s propensity score, representing the probability of disease development conditional on observed covariates. The stratification by propensity score allows for the adjustment of baseline confounders and ensures that the treated and control groups are comparable in terms of their likelihood of developing the disease. The p-value derived from the CoxPH model indicates whether the treated cohort has a statistically significant lower risk of developing the disease compared to the control cohort within the observation period, after adjusting for the propensity score. Kaplan-Meier (KM) curves were fitted to compare the disease-free survival probabilities over time between the treated and control cohorts.

### In-vitro validation of potential repurposed drugs for prostate cancer

The prostate cancer cell lines C4-2B, DU145, and PC3 were obtained from American Type Culture Collection (ATCC). Taxane-resistant DU145 (DU145-TxR) and taxane-resistant PC3 (PC3-TxR) were generated via paclitaxel exposure^91^. All cell lines were cultured in RPMI medium containing 10% FBS and penicillin-streptomycin (15140-122, Invitrogen). Cells were cultured at 37°C in 5% CO2. Candidate repurposed drugs, including bortezomib (S1013) and vorinostat (S1047) were purchased from Selleck Chemicals. The cell proliferation reagent WST-1 (11644807001) was purchased from Roche for cytotoxicity assessment.

Each STDrug repurposed drug was diluted in DMSO and added to each well so that the DMSO concentration in the medium was equal and less than 1%. Cells were seeded at a density of 5.0×104 cells/well in 96-well cell culture plates and cultured in 100uL of medium containing multiple concentration of drugs. The effect of the drugs were evaluated by a cytotoxicity assay using WST-1 reagen. Cells were cultured in a CO_2_ incubator at 37°C. After 48 hours of incubation, 10 µl of WST-1 reagent was added to 100 µl of medium in each well, and the plates were incubated at 37°C in a CO_2_ incubator for 1-2 hours. Using a Synergy H1 multi-detection reader (BioTek), the background absorption

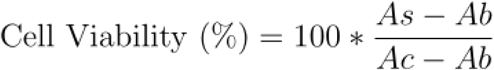

at a wavelength of 650 nm was measured and subtracted from the actual absorption value at a wavelength of 450 nm. Absorbance was measured for the drug-exposed group, negative control group, and blank group. Cell viability was calculated as where As is absorbance of sample, Ab is absorbance of blank (culture medium without cells), and Ac is absorbance of blank (culture medium without cells). The IC50 was then calculated using GraphPad Prism^92^.

## Acknowledgments

The authors acknowledge all lab members of Garmire Group for helpful discussions.

## Author contributions statement

L.X.G conceived this project and supervised the study. A.K. and S.Z. developed the first version of the tool. Y.Y. and T.U. modified the pipeline, carried the analysis, and wrote the manuscript. S.K., supervised by E.K, performed in-vitro drug validation in prostate cancer cell lines. X.Y. performed in-silico drug validation. Y.D. assisted the functional downstream analysis. T.C. assisted the disease-control balanced clustering implementation. H.L assisted the data simulation. X.G. assisted with technical troubleshooting and packaging STDrug. All authors have read and approved the final manuscript.

## Data availability

All pre-processed data will be made publicly available upon the publication of this manuscript.

